# Lower brain pH as a shared endophenotype of psychotic disorders

**DOI:** 10.1101/083550

**Authors:** Hideo Hagihara, Vibeke S Catts, Yuta Katayama, Tsuyoshi Takagi, Freesia L Huang, Kuo-Ping Huang, Shunsuke Ishii, Isabella A Graef, Gerald R Crabtree, Keiichi I Nakayama, Cynthia Shannon Weickert, Tsuyoshi Miyakawa

## Abstract

Lower pH is a well-replicated finding in the postmortem brains of patients with schizophrenia and bipolar disorder. Interpretation of the data, however, is controversial as to whether this finding reflects a primary feature of the diseases or is a result of confounding factors such as medication, postmortem interval, and agonal state. To date, systematic investigation of brain pH has not been undertaken using animal models, which can be studied without confounds inherent in human studies. In the present study, we first confirmed that the brains of patients with schizophrenia and bipolar disorder exhibit lower pH values by conducting a meta-analysis of existing datasets. We then utilized neurodevelopmental mouse models of psychiatric disorders in order to test the hypothesis that lower brain pH exists in these brains compared to controls due to the underlying pathophysiology of the disorders. We measured pH, lactate levels, and related metabolite levels in brain homogenates from three mouse models of schizophrenia (*Schnurri-2* KO, forebrain-specific *calcineurin* KO, and *neurogranin* KO mice) and one of bipolar disorder (*Camk2a* HKO mice), and one of autism spectrum disorders (*Chd8* HKO mice). All mice were drug-naïve with the same postmortem interval and agonal state at death. Upon postmortem examination, we observed significantly lower pH and higher lactate levels in the brains of model mice relative to controls. There was a significant negative correlation between pH and lactate levels. These results suggest that lower pH associated with increased lactate levels is a pathophysiology of such diseases rather than mere artifacts.

## Introduction

Schizophrenia, bipolar disorder, and autism spectrum disorders (ASD) are highly heritable psychiatric conditions, with clinical features transcending diagnostic categories^1,2^. Accumulating evidence indicates that some genetic influences^3–6^, gene expression abnormalities^7,8^, and neuronal dysfunctions^9,10^ associated with these conditions overlap, suggesting a common underlying biological basis. However, the shared neurobiological alterations among the three conditions remain largely unknown.

A number of postmortem studies have indicated that pH is lower in the brains of patients with schizophrenia and bipolar disorder^8,11–19^. Lower brain pH has also been observed in patients with ASD^20^. In general, pH balance is considered critical for maintaining optimal health, and low pH has been associated with a number of somatic disorders^21–23^. Therefore, it is reasonable to assume that lower pH may exert a negative impact on brain function and play a key role in the pathogenesis of various psychiatric disorders. However, lower brain pH has largely been considered as an artifact^11,24–27^ rather than a pathophysiology of such disorders^13,28^ for two main reasons. One is that chronic treatment with antipsychotics may affect brain pH by increasing lactate levels in rats^11^, and most patients with these disorders receive chronic antipsychotics treatment throughout their lives. Another is that the agonistic state experienced before death decreases brain pH^25–27^ and this state could be different in patients with psychiatric disorders in comparison to controls. In human postmortem studies, it is technically difficult to exclude such confounding factors and to determine whether lower pH and increased lactate levels are indeed artifacts.

In the present study, we first confirmed that patients with schizophrenia and bipolar disorder exhibit lower postmortem brain pH by conducting a meta-analysis of publicly available datasets. We then measured brain pH in multiple mouse models of psychiatric disorders, which are devoid of such confounding factors, in order to test the hypothesis that lower brain pH is a pathophysiology or an endophenotype rather than an artifact in a subgroup of psychiatric disorders. We also measured lactate levels, increases in which have frequently been linked to lower pH in the brains of patients with psychiatric disorders^11,13,29^. To our knowledge, the present study is the first to systematically evaluate pH and lactate levels in mouse models of psychiatric disorders which eliminate the confounds inherent in the human studies.

For the mouse models of psychiatric disorders, we focused on the ones reported to have neurodevelopmental abnormalities in the brain, a part of which stay at pseudo-immature status^30–33^. Specifically, we measured pH, lactate, and related metabolite levels in the postmortem brains of the following mouse models: *schnurri-2* (*Shn2*) knockout (KO) mice^34^, forebrain-specific *calcineurin* (Cn) KO mice^35–38^ and *neurogranin* (*Nrgn*) KO mice^39–41^ as a model of schizophrenia; mice with heterozygous knockout of the calcium/calmodulin-dependent protein kinase II alpha (*Camk2a* HKO mice)^42,43^ as a model of bipolar disorder;, and mice with heterozygous knockout of the long isoform of chromodomain helicase DNA-binding protein 8 (*Chd8* HKO mice)^33^ as a model of ASD. These mouse strains have mutations in the genes implicated in the respective disorders and exhibit molecular and behavioral abnormalities relevant to each condition, indicating good construct and face validities, respectively (as described in detail in Materials and Methods).

## Materials and Methods

### Human data

Nine publicly available datasets were utilized in the present study (Supplementary Table 1): four schizophrenia datasets (GSE17612, GSE21935, GSE21138; NSWBTRC-SC^44^), one bipolar disorder dataset (GSE5392), and three combined schizophrenia and bipolar disorder datasets (Stanley Medical Research Institute [SMRI] Collection A, SMRI Collection C, GSE35977, GSE53987). We obtained data regarding postmortem interval and age from these studies and data regarding medication from SMRI Collection A and SMRI Collection C.

### Animals

We measured pH, lactate, and related metabolite levels in *Shn2* KO mice^34^ (n = 5, 6 [controls, mutants]), *Cn* KO mice^35–38^ (n = 6, 5), *Nrgn* KO mice^39–41^ (n = 6, 5), *Camk2a* HKO mice^42,43^ (n = 5, 5) and *Chd8* HKO mice^33^ (n = 5, 5), and their corresponding control mice. Both male and female mice were used in the present study, as no difference in pH between genders has been observed^45^. All mice were between 19 and 45 weeks of age, and no significant difference in age was observed between controls and mutants within each strain. All animal experiments were approved by the Institutional Animal Care and Use Committee of Fujita Health University, based on the Law for the Humane Treatment and Management of Animals and the Standards Relating to the Care and Management of Laboratory Animals and Relief of Pain. Every effort was made to minimize the number of animals used.

Shn2 was originally identified as a nuclear factor-кB (NF-кB) site-binding protein that tightly binds to the enhancers of major histocompatibility complex (MHC) class I genes and acts as an endogenous inhibitor of NF-κB.^46^ Its deficiencies in Shn2 may cause mild chronic inflammation in the brain and confer molecular, neuronal, and behavioral phenotypes relevant to schizophrenia in mice^34^. Genome-wide association studies (GWASs) have identified a number of single nucleotide polymorphisms (SNPs) in the MHC region associated with schizophrenia^47–49^. *Shn2* KO mice exhibit multiple abnormal behaviors related to schizophrenia, including increased locomotor activity, deficits in working memory, abnormal social behavior and impaired prepulse inhibition, which are commonly observed in *Cn* KO mice^35,36^ and *Nrgn* KO mice^39–41^ as well.

Calcineurin (Cn) is a calcium-dependent protein phosphatase and has been implicated in synaptic plasticity^50^. *CN* has been reported to be associated with schizophrenia^51–53^, and altered expression of calcineurin has been observed in the postmortem brains of patients with schizophrenia^54,55^. Forebrain-specific *Cn* KO mice exhibit behavioral and cognitive abnormalities related to schizophrenia^35,36^. Deficits in synaptic transmission in the frontal cortex have been suggested to be the underlying mechanism of working memory impairment in these mice^38^. In addition, *Cn* KO mice exhibit disruption in ripple-associated information processing in the hippocampal CA1, which is implicated in cognitive impairments associated with schizophrenia^37^.

Neurogranin (Nrgn) is a calmodulin-binding protein that modulates activity of the Camk2 protein downstream of *N*-methyl-d-aspartic acid (NMDA) receptors, and is implicated in synaptic plasticity^39^. GWAS revealed significant association with SNPs located upstream of the *NRGN*^49^, a finding recently confirmed by a large-scale GWAS^56^, strongly suggesting that *NRGN* is a susceptibility gene for schizophrenia. *Nrgn* KO mice exhibit behavioral phenotypes related to schizophrenia^39–41^.

Camk2 is a major downstream molecule of the NMDA receptor and is thought to play an essential role in synaptic plasticity. A recent study demonstrated genetic association of *CAMK2A* with bipolar disorder^57^, and decreased mRNA expression has been observed in the frontal cortex of patients with bipolar disorder^58^. In addition, the *Camk2a* gene was identified as one of the top candidate genes for bipolar disorder by a meta-analysis that integrated genetic and genomic data from both human and animal studies^59^. At cellular level, neuronal hyperexcitability, which we previously detected in the hippocampal granule cells of *Camk2a* HKO mice^42^, was also found in the granule cell-like neurons differentiated from induced pluripotent stem cells (iPSCs) derived from patients with bipolar disorder^60^. *Camk2a* HKO mice exhibit abnormal behaviors, such as deficits in social activity and working memory, which are analogous to those in patients with bipolar disorder/schizophrenia^42^. In addition, these mutant mice exhibit infradian cyclic activity levels, which may reflect infradian oscillation of mood substantially observed in patients with bipolar disorder^43^. These findings suggest that *Camk2a* HKO mice have construct and face validity as a model of psychiatric disorders, especially of bipolar disorder.

Chd8, a member of the chromodomain helicase DNA-binding family of proteins, is known to act as a chromatin-remodeling factor. Recent exome sequencing analyses have identified a number of *de novo* mutations in a variety of genes in individuals with ASD, further revealing that *CHD8* is the most frequently affected gene^61–64^. *Chd8* HKO mice exhibit behavioral abnormalities reminiscent of ASD in humans, including increased anxiety, increased persistence and abnormal social interaction^33^. *Chd8* deficiency induces aberrant activation of RE1 silencing transcription factor (REST), a molecular brake of neuronal development, resulting in neurodevelopment abnormalities in mice^33^.

Collectively, these findings indicate that the mouse models used in the present study exhibit good construct and face validities for their respective disorders.

### Measurement of pH

Mice were sacrificed by cervical dislocation followed by decapitation, following which whole brains were removed. The brains were immediately frozen in liquid nitrogen and stored at −80°C until use. We measured brain pH basically as previously described^11^. Briefly, the brains were homogenized using the tissue homogenizer attached with a conical pestle in ice-cold distilled H_2_O (5 mL per 500 mg of tissue). The pH was measured using a pH meter (LAQUA F-72, Horiba Scientific, Kyoto, Japan) after a three-point calibration at pH 4.0, pH 7.0 and pH 9.0. The pH of the samples from control and mutant mice were read in triplicate for each sample. After pH measurement, homogenates were immediately frozen and stored at −80°C until required for further analyses.

### Lactate and glucose measurements

The concentration of lactate in the brain homogenates was determined using a multi-assay analyzer (GM7 MicroStat; Analox Instruments, London, UK) according to manufacturer’s instructions. In our prior tests using several samples, we loaded 5 μl, 10 μl and 20 μl of supernatants to the instrument, observing that the measurements increased linearly in a volume-dependent manner (r^2^ > 0.99). Based on these results, we used 20 μl of supernatants for each sample for lactate measurements. Likewise, glucose concentrations in 20 μl supernatant samples were determined using a multi-assay analyzer following calibration with 10 mmol/ml glucose standard solution. To normalize the effects of differences among strains, such as genetic background and age, *z*-scores for pH and lactate levels were calculated within each strain and used for the correlation analysis.

### Pyruvate measurement

Pyruvate concentrations in 20 μl supernatant samples were determined using a pyruvate assay kit (BioVision, Mountain View, CA, USA). The fluorescence intensities were measured using a microplate reader equipped with a spectrofluorometer (ARVO X, PerkinElmer).

### Adenosine diphosphate/adenosine triphosphate (ADP/ATP) ratio

An ADP/ATP Ratio Assay Kit (BioVision) was used to measure the ADP and ATP concentrations according to the manufacturer’s instructions.

### Bioinformatics analysis of transcriptome data

We used the following mouse brain transcriptome data: frontal cortex and hippocampal dentate gyrus of *Shn2* KO mice (microarray)^34^, hippocampal dentate gyrus of *Camk2a* HKO mice (microarray)^65^, and whole brains of *Chd8* HKO mice (RNA-sequencing)^33^. Gene expression patterns of the frontal cortex of *Camk2a* HKO mice (n = 6, 6) and hippocampal DG of *Cn* KO mice (n = 6, 6) were analyzed via microarray (Mouse Genome 430 2.0 Array; Affymetrix, Santa Clara, CA, USA), as previously described^34^. Gene expression patterns of the frontal cortex and hippocampal DG of *Nrgn* KO mice (n = 5, 5) were analyzed via RNA-sequencing using the HiSeq platform basically according to the manufacturer’s instructions (Illumina, San Diego, CA, USA). Genes with an absolute fold change > 1.2 and a *t*-test *P*-value < 0.05 (mutants vs. controls; without correction for multiple testing) were imported into the bioinformatics tool BaseSpace (Illumina), with which the gene expression data obtained from different platforms can be matched. Genes with altered expression in at least four out of the eight datasets (yielding 80 features; Supplementary Table 2) were selected based on the criteria of the BaseSpace tool and assessed for enrichment in biological themes using the DAVID functional annotation clustering tool, ADGO, and GOToolBox, in which the default feature listings and algorithm settings were used.

## Results

### Meta-analysis of human brain pH studies

We first re-evaluated the results of postmortem studies of brain pH in patients with schizophrenia and bipolar disorder by conducting a meta-analysis of publicly available datasets. We searched the National Center for Biotechnology Information Gene Expression Omnibus (NCBI GEO), ArrayExpress, and Stanley Medical Research Institute (SMRI) databases, and found nine studies that included individual brain pH data from patients with schizophrenia, bipolar disorder, or both as well as from healthy control participants (Supplementary Table 1). A two-way analysis of variance (ANOVA) revealed a significant effect of condition (F_2_,_645_ = 3.35, *P* = 3.09 × 10^−10^) and study (F_8_,_645_ = 10.00, *P* = 2.00 × 10^−16^) as well as between the two factors (F_11_,_645_ = 47.66, *P* = 0.043) (Figure 1). *Post hoc* comparisons with Tukey’s honest significant difference test indicated a lower brain pH in both patients with schizophrenia (*P* < 1.0 × 10^−7^) and bipolar disorder (*P* = 0.00036) compared to healthy controls, and no significant difference between the two conditions (*P* = 0.56). The results of our meta-analysis therefore support the finding of lower brain pH in patients with schizophrenia and bipolar disorder.

**Figure 1.**
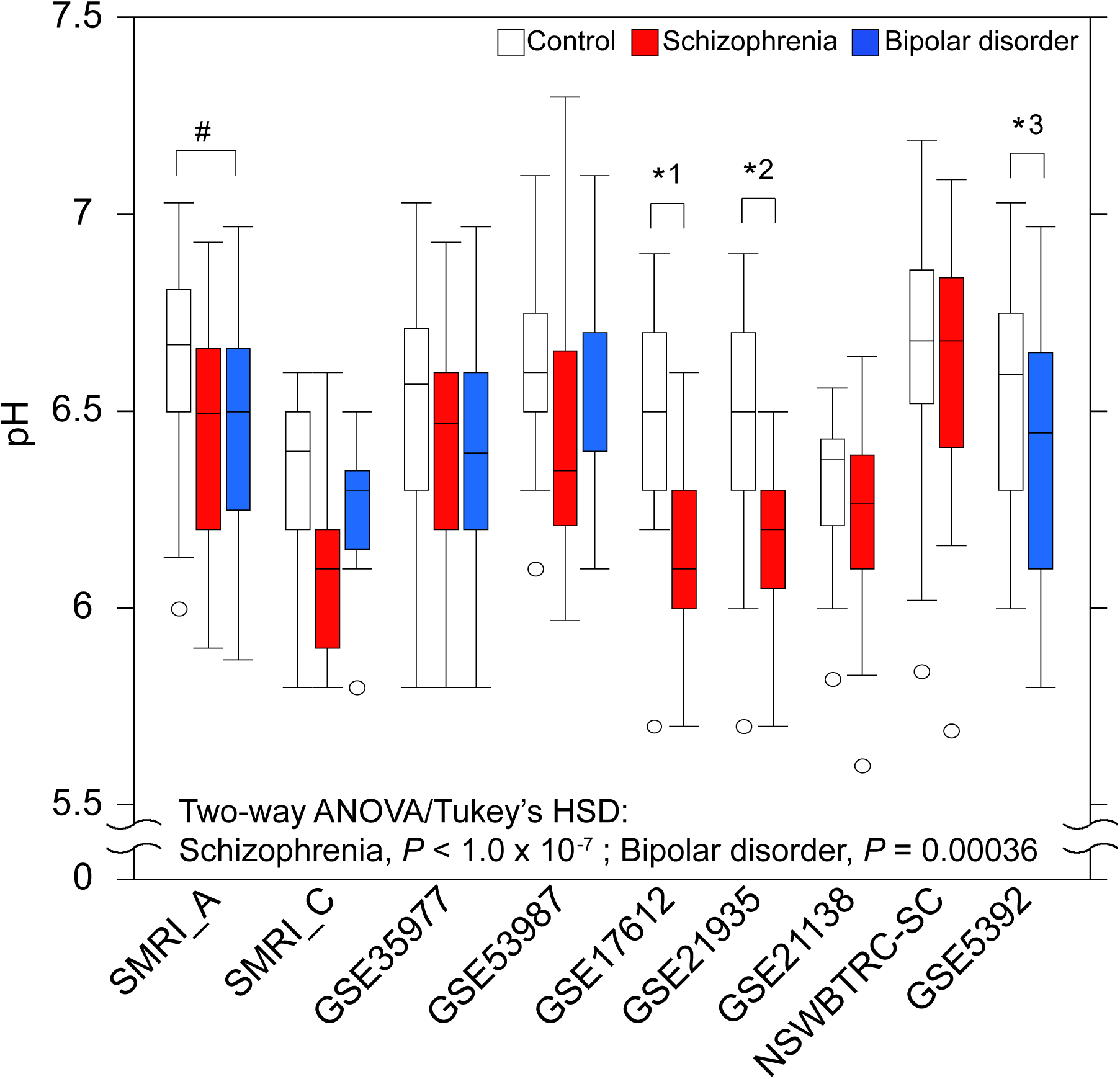
Lower pH in the postmortem brains of patients with schizophrenia and bipolar disorder revealed by meta-analysis of publicly available data. Box plot of pH in the brain of control participants (*white box*), patients with schizophrenia (*red box*) and patients with bipolar disorder (*blue box*). *#P* = 0.017; One-way analysis of variance (ANOVA)/Tukey’s honest significant difference test. *^1^ *P* = 8.0 × 10^−6^, *^2^ *P* = 1.3 × 10^−4^, *2 *P* = 0.027; Student’s *t*-test. The boxes represent the interquartile range between first and third quartiles, the whiskers the maximum and minimum values and the circles population outliers.

Brain pH was not correlated with lifetime use of antipsychotics (measured as fluphenazine equivalents) in a schizophrenia group (Pearson’s *r* = −0.27, *p* = 0.12), a bipolar disorder group (*r* = −0.27, *P* = 0.13), or a group including both conditions (*r* = −0.15, *P* = 0.23) from the SMRI Collection A cohort (Supplementary Figure 1a). No correlation between pH and antipsychotics was replicated in a separate cohort from SMRI Collection C (schizophrenia: *r* = 0.16, *P* = 0.58; bipolar disorder: *r* = 0.036, *P* = 0.90; both: *r* = 0.090, *P* = 0.64) (Supplementary Figure 1b). These suggest that antipsychotics treatment may not affect pH in the postmortem brains of patients with schizophrenia and bipolar disorder. On the other hand, in the combined data, pH was positively correlated with postmortem interval (*r* = 0.13, *P* = 0.0010; Supplementary Figure 1c) and negatively correlated with age (*r* = −0.13, *P* = 0.00092; Supplementary Figure 1d), suggesting that these factors may contribute to the changes in the pH of postmortem brains.

### Lower pH and increased lactate levels in the postmortem brain of mouse models of schizophrenia, bipolar disorder, and ASD

The confounding factors identified in previous studies^11,26^ are beyond investigator’s control in human postmortem brain studies. We therefore measured pH and lactate levels in the brains of mouse models of schizophrenia (*Shn2* KO, *Cn* KO, *Nrgn* KO mice), bipolar disorder (*Camk2a* HKO mice), and ASD (*Chd8* HKO mice). All the mice used were drug-naïve and sacrificed by cervical dislocation (controling for agonal state differences). The removed brains were snap-frozen within a few minutes (controling for postmortem interval differences). Brain pH was significantly lower in all five mutant strains examined relative to the corresponding controls (*Shn2* KO, 7.17 ± 0.0060, controls [Con], 7.20 ± 0.056, *P* = 0.0083; *Cn* KO, 7.08 ± 0.0057, Con, 7.13 ± 0.0080, *P* = 0.0014; *Nrgn* KO, 7.10 ± 0.017, Con, 7.16 ± 0.0080, *P* = 0.0090; *Camk2a* HKO, 7.14 ± 0.0093, Con, 7.21 ± 0.0090, *P* = 0.0014; *Chd8* HKO, 7.08 ± 0.0066, Con, 7.12 ± 0.0031, *P* = 0.00080) (Figure 2a).

**Figure 2.**
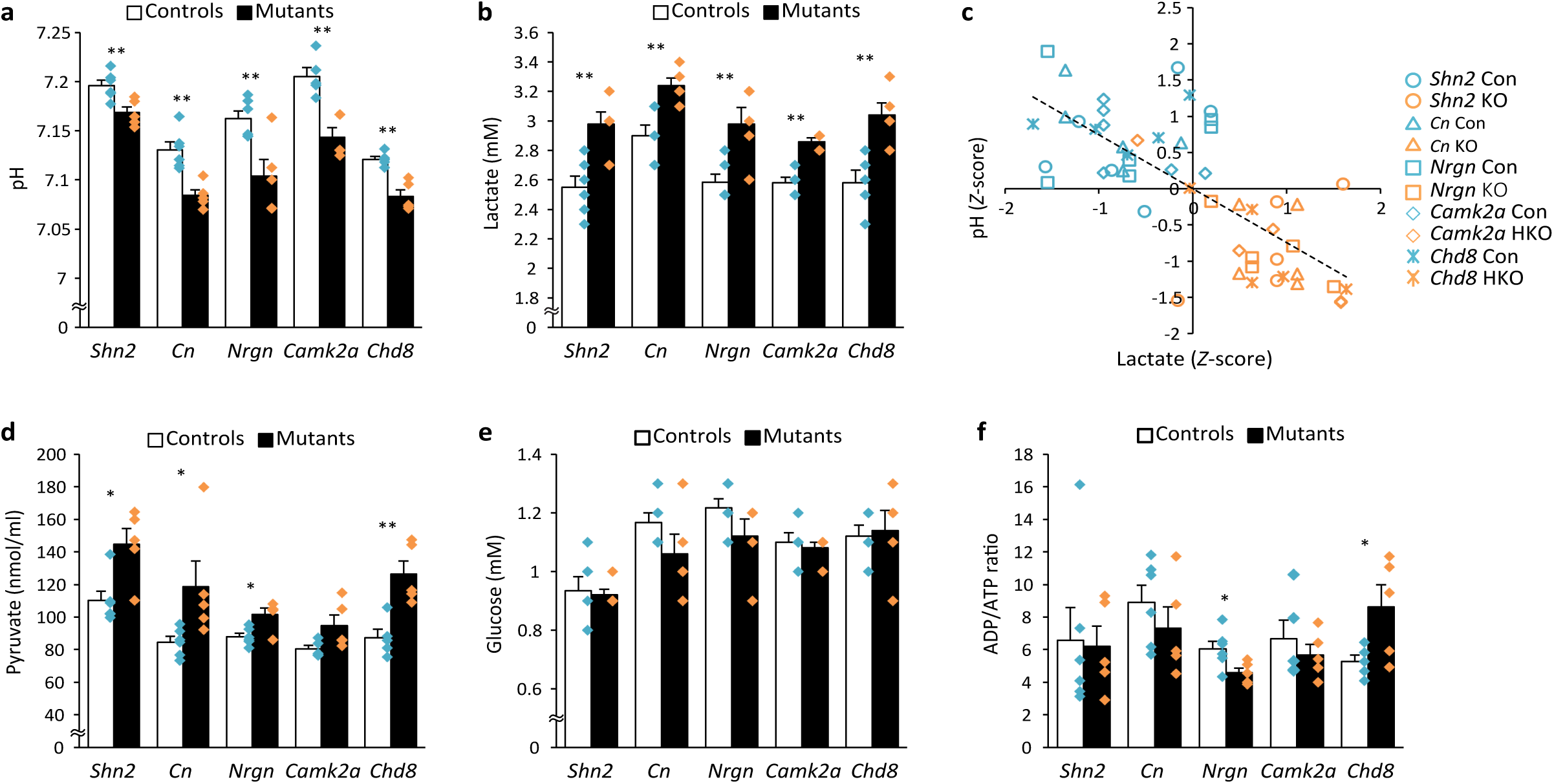
Negative correlation between lower pH and increased lactate levels in the postmortem brains of mouse models of psychiatric disorders. Bar graphs of pH (**a**), lactate levels (**b**), pyruvate levels (**d**), glucose levels (**e**), and ADT/ATP ratio (**f**) in the brains of *Shn2* KO, *Cn* KO, *Nrgn* KO, *Camk2a* HKO, and *Chd8* HKO mice and their corresponding controls (average ± SEM). Each plot represents individual mouse values. (**c**) Scatter plot showing correlations between pH and lactate levels in the mouse brain. \**P* < 0.05, \*\**P* < 0.01; Student’s *t*-test. SEM: standard error of the mean. ADP: adenosine diphosphate; ATP: adenosine triphosphate.

Significantly higher levels of lactate were observed in the postmortem brains of all mutant mice strains compared to corresponding controls (*Shn2* KO, 2.98 ± 0.080 mM, Con, 2.55 ± 0.076 mM, *P* = 0.0038; *Cn* KO, 3.24 ± 0.051 mM, Con, 2.90 ± 0.073 mM, *P* = 0.0052; *Nrgn* KO, 2.98 ± 0.11 mM, Con, 2.58 ± 0.054 mM, *P* = 0.0080; *Camk2a* HKO, 2.86 ± 0.024 mM, Con, 2.58 ± 0.037 mM, *P* = 0.00024; *Chd8* HKO, 3.04 ± 0.081 mM, Con, 2.58 ± 0.086 mM, *P* = 0.0046; Figure 2b). Analysis of the combined data expressed as the *z*-score revealed that pH was significantly negatively correlated with lactate levels (Pearson’s *r* = −0.65, *P* = 1.19 × 10^−7^; Figure 2c).

Lactate is formed from pyruvate during glycolysis. We therefore measured pyruvate levels in mutant mouse brains and observed that levels were significantly increased in *Shn2* KO (*P* = 0.011), *Cn* KO (*P* = 0.046), *Nrgn* KO (*P* = 0.011) and *Chd8* HKO mice (*P* = 0.0036) and showed increased tendency in *Camk2a* HKO mice (*P* = 0.068) (Figure 2d). Glucose levels remained unchanged in mutant mice relative to controls (Figure 2e), suggesting glucose supply/demand ratio in the brain may be comparable in these mouse models. The ADP/ATP ratio was decreased in *Nrgn* KO mice (*P* = 0.035) and increased in *Chd8* HKO mice (*P* = 0.047) (Figure 2f), suggesting a contrasting energy consumption ratio in mouse models of schizophrenia and ASD.

We then analyzed transcriptome data (Supplementary Table 2) in order to investigate the potential underlying molecular mechanisms of increased lactate levels in mutant mouse brains. The transcriptome data from five mouse strains revealed an enrichment in Wnt- and epidermal growth factor (EGF)-related pathways when analyzed with DAVID software (Supplementary Table 3). Enrichment in Wnt-related pathways was replicated in the analyses using other bioinformatics tools (ADGO and GOToolBox) using different statistical methods (Supplementary Table 3).

Since lactate is produced via glycolytic pathways in astrocytes in the brain^66^, we analyzed the transcriptome data of mutant mice with particular focus on glycolysis-related genes (Gene Ontology Consortium database), as well as those related to pyruvate metabolism. The results of the targeted gene expression analyses suggest that elevated glycolysis and pyruvate metabolism shifting toward lactate synthesis occurs in the brains of mutant mice, especially in *Shn2* KO and *Camk2a* HKO mice (Supplementary Table 4; Supplementary Figure 2).

## Discussion

In the present study, we confirmed lower pH in the postmortem brains of patients with schizophrenia and bipolar disorder by conducting a meta-analysis of existing datasets. Lower pH was also observed in five different mouse models of psychiatric disorders, all of which were drug-naïve and were controlled for other confounding factors, such as agonal state and postmortem interval. We also observed increased lactate levels in the brains of mutant mice, as well as a highly significant negative correlation between pH and lactate levels, which is consistent with the findings of previous human postmortem studies^11^. These results suggest that lower pH and increased lactate levels represent components of the underlying pathophysiology of the diseases rather than mere artifacts.

Researches have revealed that brain acidosis influences a number of brain functions, such as anxiety, mood, and cognition^67^. Acidosis may affect the structure and function of several types of brain cells,including the electrophysiological functioning of GABAergic neurons^68^ and morphological properties of oligodendrocytes^69^. Alterations in these types of cells have been well-documented in the brains of patients with schizophrenia, bipolar disorder, and ASD^70,71^ and may underlie some of the cognitive deficits associated with these disorders. Deficits in GABAergic neurons and oligodendrocytes have been identified in the mouse models of the disorders, including *Shn2* KO mice^30,34^. Brain acidosis may therefore be associated with deficits in such cell types in schizophrenia, bipolar disorder, and ASD.

A previous study indicated that chronic treatment with antipsychotics increases lactate levels in the rat cerebral cortex^11^, suggesting that such increases may be medication-related. The authors of the report, however, found no significant correlation between lactate levels and history of antipsychotic use (which was represented by chlorpromazine equivalents) in the postmortem brains of patients with schizophrenia^11^. In addition, increased lactate levels have been observed in the anterior cingulate of medication-free patients with bipolar disorder in *in vivo* spectroscopic imaging studies^72^. Furthermore, studies utilizing animal models of psychiatric disorders—including the current study—have identified increased lactate levels in mutant mouse brains^73^. In addition, increased lactate levels were associated with lower pH in the brains of mutant mice, consistent with findings from previous studies on patients with schizophrenia^11,13^. Lower brain pH has also been observed in the medication-free patients with bipolar disorder^28^. Although it remains possible that antipsychotic treatment increases lactate levels and lowers pH in the brain, the aforementioned findings suggest that such changes may occur as primary features of schizophrenia and bipolar disorder.

Interestingly, we observed that Wnt- and EGF-related pathways, which are highly implicated in somatic and brain cancers^74^, are enriched in the genes whose expressions were altered among the five mutant mouse strains. It is known that cancer cells display high rates of glycolysis, resulting in high lactate and pyruvate levels, even in normoxia^75^; this phenomenon has been referred to as the Warburg effect. Genes whose expression is known to positively regulate the Warburg effect, such as *Hk2*^76^, *Hif1a*^75^, and *Pfkfb3*^77^, were increased in the brains of any of mouse models examined in the present study, while expression of *Prkaa1*, a negative regulator of the Warburg effect^78^, was decreased (Supplementary Table 2). These findings raise the possibility that elevated glycolysis underlies the increased lactate and pyruvate levels in the brains of the mouse models of schizophrenia, bipolar disorder, and ASD. The results of the targeted gene expression analyses conducted in the present study also support the hypothesis. Glycolysis is also stimulated by the uptake of glutamate in astrocytes following neuronal excitation^79^. Dysregulation of the excitation-inhibition balance has been proposed as a candidate cause of schizophrenia, bipolar disorder, and ASD^80,81^. A shift in the balance towards excitation would result in increased energy expenditure and may lead to increased glycolysis. Indeed, *Shn2* KO mice exhibit higher glutamate levels in the hippocampus^34^. *In vivo* metabolite measurements have indicated that increased glycolysis occurs in the brains of patients with bipolar disorder^29,72^, while gene ontology analysis of microarray data has indicated that decreased glycolysis occurs in the brains of patients with schizophrenia^13^. Further studies are required to determine whether altered glycolysis rate is associated with increased lactate levels.

It has been indicted that lactate levels in the mouse brain rapidly increase after at least 1 min of decapitation as compared to *in vivo* fixation by focused microwave irradiation, which is regarded as a consequence of enhanced glycolysis under oxygen-deprived conditions^82^. While the current findings may differ from those obtained under physiological conditions, they may reflect functional changes, such as the activation of astrocytes^34,40^, which represent the main source of lactate production in the brain.

Brain pH is associated with notable changes in gene expression^16,26,45,83^ and has hence been considered as a confound for investigating changes in gene expression related to the pathophysiology of psychiatric disorders. Therefore, substantial effort has been made to match the tissue pH between patients and controls. Given that lower brain pH is a pathophysiology of certain conditions, pH-dependent changes in gene expression would not be negligible when attempting to elucidate the molecular basis of the conditions. It has been known that gene expression patterns are partially similar across diseases such as schizophrenia, bipolar disorder, and ASD^7,8^. Lower pH may underlie the similarities of gene expression patterns. Thus, pH may be an important factor in the elucidation of molecular alternations in the brains of patients with these psychiatric conditions.

## Acknowledgments

We thank Wakako Hasegawa, Yumiko Mobayashi, Misako Murai, Tamaki Murakami, Miwa Takeuchi, Satoko Hattori and Aki Miyakawa, Fujita Health University, for their technical support in this study, and Yuki Sugiura, Keio University, for helpful discussion. This work was supported by JSPS Grant-in-Aid for Scientific Research on Innovative Areas Grant Number 25116526, 15H01297, JSPS KAKENHI Grant Number 25242078, and AMED Strategic Research Program for Brain Sciences.

## Author contributions

Conceived and designed the experiments: HH and TM. Contributed materials: VC, YK, TT, FH, KH, SI, IG, GC, KN and CW. Analyzed the data: HH and TM.

## Conflict of interest

Dr. Tsuyoshi Miyakawa and Dr. Cynthia Shannon Weickert both receive research grants from Astellas Pharma Inc. Dr. Cynthia Shannon Weickert is a consultant for Lundbeck, Australia Pty Ltd. Other authors have no conflict of interests to declare.

## Supplementary information

**Supplementary Figure 1. Correlations between pH and lifetime antipsychotic use, postmortem interval, and age** Scatter plots showing correlations between pH and lifetime antipsychotic (fluphenazine equivalents) use in the SMRI collection A (**a**) and SMRI collection C (**b**) datasets. Scatter plots showing the correlation between pH and postmortem interval (controls: *r* = 0.027, *P* = 0.66; schizophrenia: *r* = 0.27, *P* = 2.1 × 10^−5^; bipolar disorder: *r* = 0.14, *P* = 0.085; Total: *r* = 0.13, *P* = 0.0010) (**c**), and age (controls: *r* = −0.14, *P* = 0.021; schizophrenia: *r* = −0.22, *P* = 0.00075; bipolar disorder: *r* = 0.14, *P* = 0.096; Total: *r* = −0.13, *P* = 0.00092) (**d**). SMRI: Stanley Medical Research Institute.

**Supplementary Figure 2. Potentially elevated glycolysis in the brains of mouse models of psychiatric disorders** Glycolysis-related genes whose expression was altered in the brains of mouse models of psychiatric disorders were mapped in a schematic of the glycolysis pathway.

**Supplementary Table 1. Patient characteristics** Antipsychotic dose (mg) is measured as fluphenazine equivalents. M, male; F, female; na, not available.

**Supplementary Table 2. Genes whose expression was altered in the brains of mouse models of psychiatric disorders** Genes whose expression was altered in at least four out of eight mouse datasets were processed for pathway analyses.

**Supplementary Table 3. Pathway analyses of the genes whose expression was altered in the brains of mouse models of the psychiatric disorders using DAVID, ADGO, and GoToolBox** The top 20 pathways (ranked based on the *P*-value) are shown for each analysis.

**Supplementary Table 4. Expression patterns of genes encoding enzymes related to glycolysis pathway in the brains of mouse models of psychiatric disorders**

